# Spatiotemporal bias of the human gaze toward hierarchical visual features during natural scene viewing

**DOI:** 10.1101/2022.06.08.495305

**Authors:** Kazuaki Akamatsu, Tomohiro Nishino, Yoichi Miyawaki

## Abstract

The human gaze is directed at various locations from moment to moment in acquiring information necessary to recognize the external environment at the fine resolution of foveal vision. Previous studies showed that the human gaze is biased to particular locations in the visual field at a particular timing, but it remains unclear what visual features produce such spatiotemporal bias. In this study, we used a deep convolutional neural network model to extract hierarchical visual features from natural scene images and evaluated how much the human gaze is attracted to the visual features in space and time. Eye movement measurement and visual feature analysis using the deep convolutional neural network model showed that the gaze was strongly attracted to spatial locations containing higher-order visual features than lower-order visual features and conventional saliency. Analysis of the gaze time course revealed that the bias to higher-order visual features was prominent within a short period after the beginning of observation of the natural scene images. These results demonstrate that higher-order visual features are a strong gaze attractor in both space and time, suggesting that the human visual system uses foveal vision resources to extract information from higher-order visual features with higher spatiotemporal priority.

## Introduction

Humans acquire visual information of the external world while moving their eyes to direct their gaze at various locations. The eye movement is essential for accurate visual recognition because the spatial resolution of human vision is not uniform over eccentricity—i.e., the foveal vision possesses visual acuity higher than that of peripheral vision (Anstis, 1974). Given this constraint, the human visual system prioritizes particular locations by directing gaze and processes the information using the foveal vision at a high spatial resolution. However, what visual features determine such gaze biases in observed images remains under debate.

Conventionally, “saliency” has been considered an influential feature, with much experimental evidence showing that the human gaze is frequently directed to spatial locations with higher saliency (Itti et al., 1998). Saliency is typically defined as compositions of lower-order visual features such as luminance contrast, color contrast, and edges (Itti et al., 1998; Walther and Koch, 2006), although there are several variants of its definition (Cerf et al., 2009). Computational models have also revealed that gazed locations can be predicted using a saliency map that quantifies a value of saliency at each spatial location in observed images (Itti et al., 1998; Walther and Koch, 2006). However, saliency requires computation at not only a local scale but multiple scales including large spatial coverage (Itti et al., 1998; Walther and Koch, 2006). It is thus difficult to infer what visual cortical area is primarily involved in multiscale feature extraction processes and the generation of gaze bias.

Recent studies showed that activation of a higher layer of deep convolutional neural network (DCNN) models can be used to predict the gaze bias to particular locations in observed images. The prediction accuracy is much higher than that of the conventional saliency map (Kümmerer et al., 2015; Huang et al., 2015 Jetly et al., 2016; Kruthiventi et al., 2017; Kümmerer et al., 2017), implying that higher-order visual features are more informative than lower-order visual features in predicting the gaze bias in space. However, in these studies, the DCNN models were combined with a multilayer neural network module whose parameters were tuned to predict the gaze bias, and the combination obscured whether the gaze was really attracted toward the spatial locations containing the higher-order visual features. In fact, gazed locations were predicted when only lower-order image information was input to the multilayer neural network module, and the accuracy was even better than that when the deep layer activation of the DCNN model was used (Kümmerer et al., 2017). Hence, it remains unclear whether the gaze is attracted to spatial locations containing higher-order visual features or the multilayer neural network module merely contributes to accurate gaze prediction.

Similar issues exist when considering what visual features attract the gaze earlier in time. Previous studies suggest that spatial locations having higher saliency attract the gaze within a short period after image viewing starts (Anderson et al., 2015; Anderson et al., 2016). However, recent studies argue that such gaze bias in time can be predicted by activation of the deep layer of DCNN models more accurately than by using the saliency-based model (Schütt et al., 2019). In this study, the gaze was predicted by attaching an additional neural network module to the DCNN model, and the contribution of higher-order visual features to the gaze bias over time thereby remains uncertain.

To resolve the above issues, we examine the direct relationship between the spatiotemporal gaze bias and the visual features by mapping gazed locations and DCNN-extracted hierarchical visual features into observed images. Here, we use a DCNN model that was pretrained using an independent dataset of natural images (Krizhevsky et al., 2011) as a hierarchical visual feature extractor without any additional modules for gaze prediction. As many studies have demonstrated, the DCNN can be considered a good model of hierarchical visual systems of the human brain, with each layer responding to visual features of different levels of complexity from low level to high level along its hierarchy. This similarity allows us to infer what visual areas or levels of visual information processing in the brain affect the gaze bias in observed images. Furthermore, because the DCNN model is used as an independently pretrained filter to extract hierarchical visual features in a bottom-up fashion, we can examine what and when visual features attract the gaze. In other words, this methodology is expected to reveal spatiotemporal characteristics of the gaze bias to visual features in observed images. Our approach is expected to provide not only systematic knowledge about feature-based human gaze attraction but also useful insight into effective visual designs.

## Results

We selected various images from large-scale natural scene databases (Mottaghi et al., 2014; Zhou et al., 2016) and used them as visual stimuli in experiments. Using these visual stimuli, we performed encoding runs in which participants observed the visual stimuli with free eye movements, followed by recognition runs in which the participants observed the visual stimuli with free movements and answered whether the observed stimuli were presented in the preceding encoding run. The recognition task accuracy was significantly higher than the chance level for all participants (Supplementary figure 2; binomial test p < 0.05 for each participant; mean ± standard deviation across participants, 91.25% ± 6.74%), indicating that all participants engaged in the experiment.

The eye movement of the participants was measured using an infrared camera system at a sampling rate of 1000 Hz, which is fast enough to capture the participants’ gazed location. The eye movement was recorded for both the encoding and recognition runs (Figure 1), but only results for the encoding runs are presented in this section and the recognition runs were not analyzed because half of the visual stimuli in the recognition runs had been presented in the encoding runs, which might affect perceptual states.

**Figure 1:**
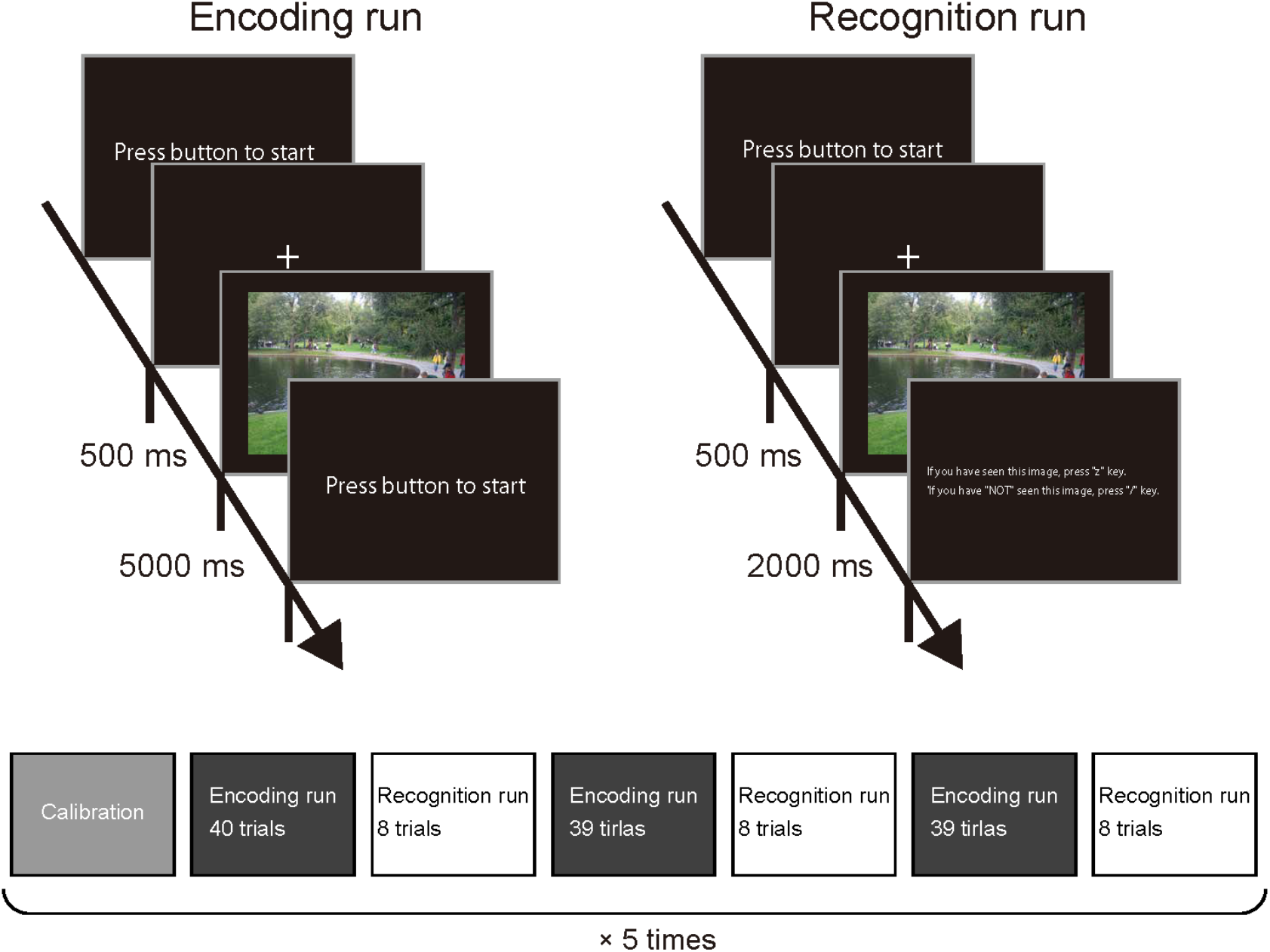
Experimental design. The experiments comprised encoding and recognitions runs. In the encoding runs, the participants were asked to observe presented visual stimuli with eye movements allowed. In the recognition runs, the participants were asked to observe presented visual stimuli with eye movements allowed and to answer whether each image had been presented in the preceding encoding runs. The encoding and recognition runs were combined and performed in the order presented in the figure, preceded by calibration of the eye tracker. This sequence was performed five times for each participant.

For each visual stimulus, we generated feature maps computed from unit activations in convolutional layers 1 to 5 of AlexNet (Krizhevsky et al., 2012), one of the most accepted DCNN models for the hierarchical human visual system (Yamins et al., 2014; Güçlü et al., 2015; Horikawa et al., 2017). The feature maps visualize spatial distributions of the intensity of visual features that contribute to unit activation in each convolutional layer using the SmoothGrad technique (Smilkov et al., 2017), which backprojects the unit activation in each layer into the image-pixel space (Figure 2A). In addition, we also calculated saliency maps (Figure 2A), which have been typically used for examining the relationship between visual features and eye movement. In this study, saliency maps were computed using SaliencyToolbox (Walther and Koch, 2006).

**Figure 2:**
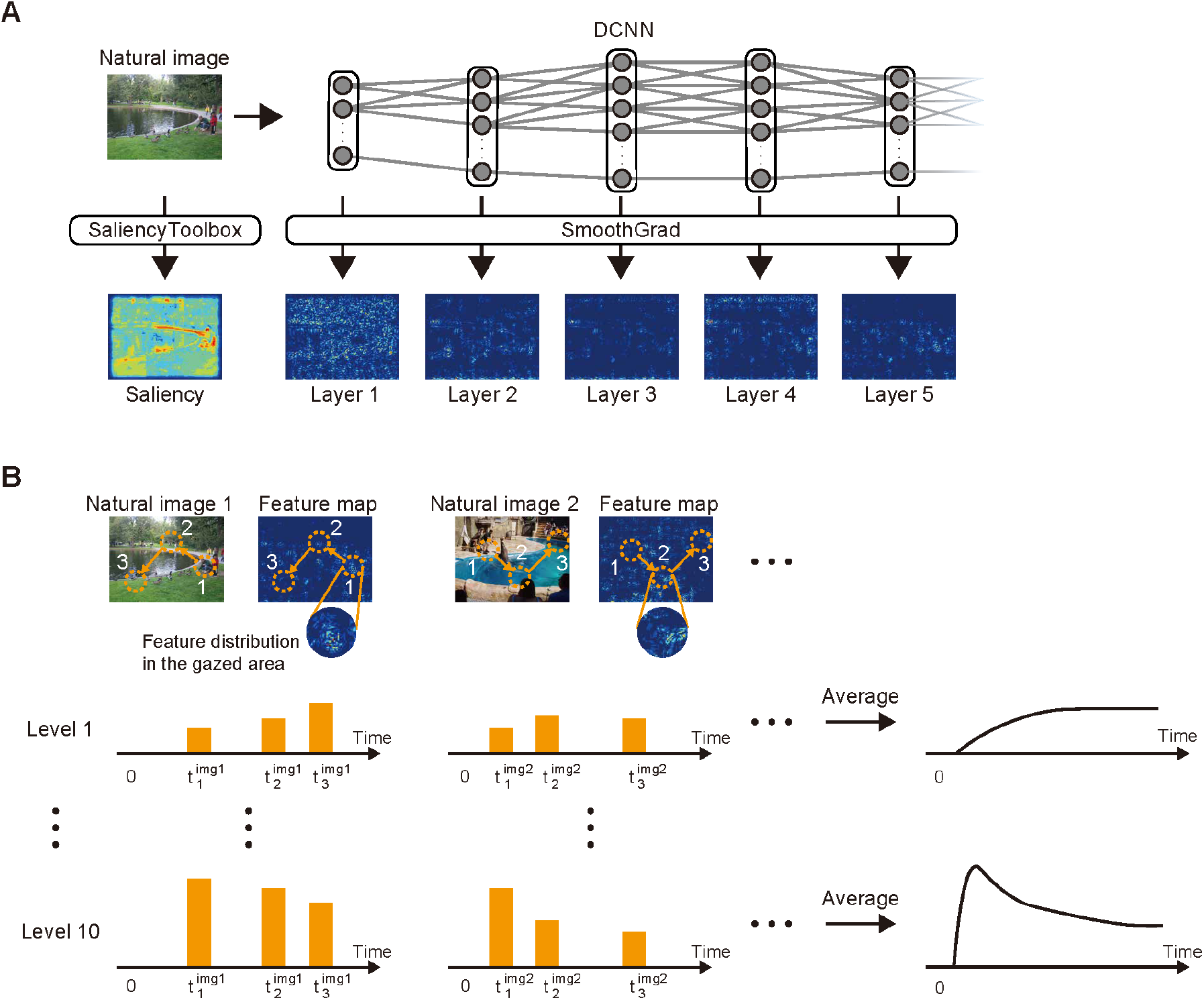
Feature maps and gaze attraction. A) Feature maps corresponding to hierarchical layers of the DCNN and saliency. The unit activation of each layer of the DCNN is backprojected to the image-pixel space using the SmoothGrad technique for each visual stimulus. The intensity of features is represented by pseudocolor (blue, low; red, high). B) Time course of the gaze attraction of visual features. The combination of gaze measurement data and the feature map provides how much of a visual feature is contained in each gazed area for a specified intensity level. This procedure defines the gaze attraction of each visual feature for the intensity level at the time that the gaze occurs. The average of these values over all visual stimuli then gives the time course of the gaze attraction of each visual feature for each intensity level.

### Gaze attraction to visual features

We used the above experimental datasets to quantify what visual features attracted the gaze at each time after the stimulus presentation (Figure 2B). The eye movement data provide a series of gazed locations, around which we can calculate how much visual features exist. We interpreted a high calculated value as a consequence of the gaze being attracted by the visual feature existing at the location. We thus refer to this value as the gaze attraction for each visual feature. Because the gaze attraction naturally depends on the intensity of the visual feature, we discretized the feature intensity into ten levels between the minimum to the maximum value in each entire stimulus image and evaluated the gaze attraction for each intensity level separately. Repeating this procedure for each gazed location at gaze timing and taking the average over all presented stimuli, we can illustrate the time course of the gaze attraction for each visual feature at different intensity levels.

### Spatial gaze bias

Figure 3 shows the representative examples of the time courses of the gaze attraction for the three intensity levels of 1 (lowest), 5 (middle), and 10 (highest), compared with the chance level calculated from the randomized gazed locations (see Methods). For the visual features at low-level intensity (top row in Figure 3), the gaze attraction was almost at the same level as chance (layers 1 to 5) or lower (saliency). There was no major difference between visual features except saliency because all feature intensities were low and their characteristics were thus missed. The difference from the chance level became evident as the intensity increased (middle to bottom rows in Figure 3), indicating that the gaze was attracted to locations with intense visual features (Figure 4; two-way analysis of variance (ANOVA), p < 0.05 for the main effect in terms of feature intensity). The difference between the visual features also became prominent for the higher intensity levels (middle to bottom rows in Figure 3). The mean gaze attraction over time, or spatial gaze bias (see Methods), progressively increased with the hierarchy of the convolutional layers (Figure 4; two-way ANOVA, Bonferroni-corrected p < 0.05 for multiple comparisons; the spatial gaze bias increased in the order of saliency, layer 1, 2, 3, 4, and 5 (pair-wise significance was not found between layer 3 and 4; all other pair-wise comparisons were significant), indicating that the higher-order visual features activating deep layers of the DCNN were stronger gaze attractors than the lower-order visual features activating shallow layers of the DCNN. These results are consistent with the results of previous studies showing deep features outperforming conventional saliency in the prediction of the fixation map (Kümmerer et al., 2017).

**Figure 3:**
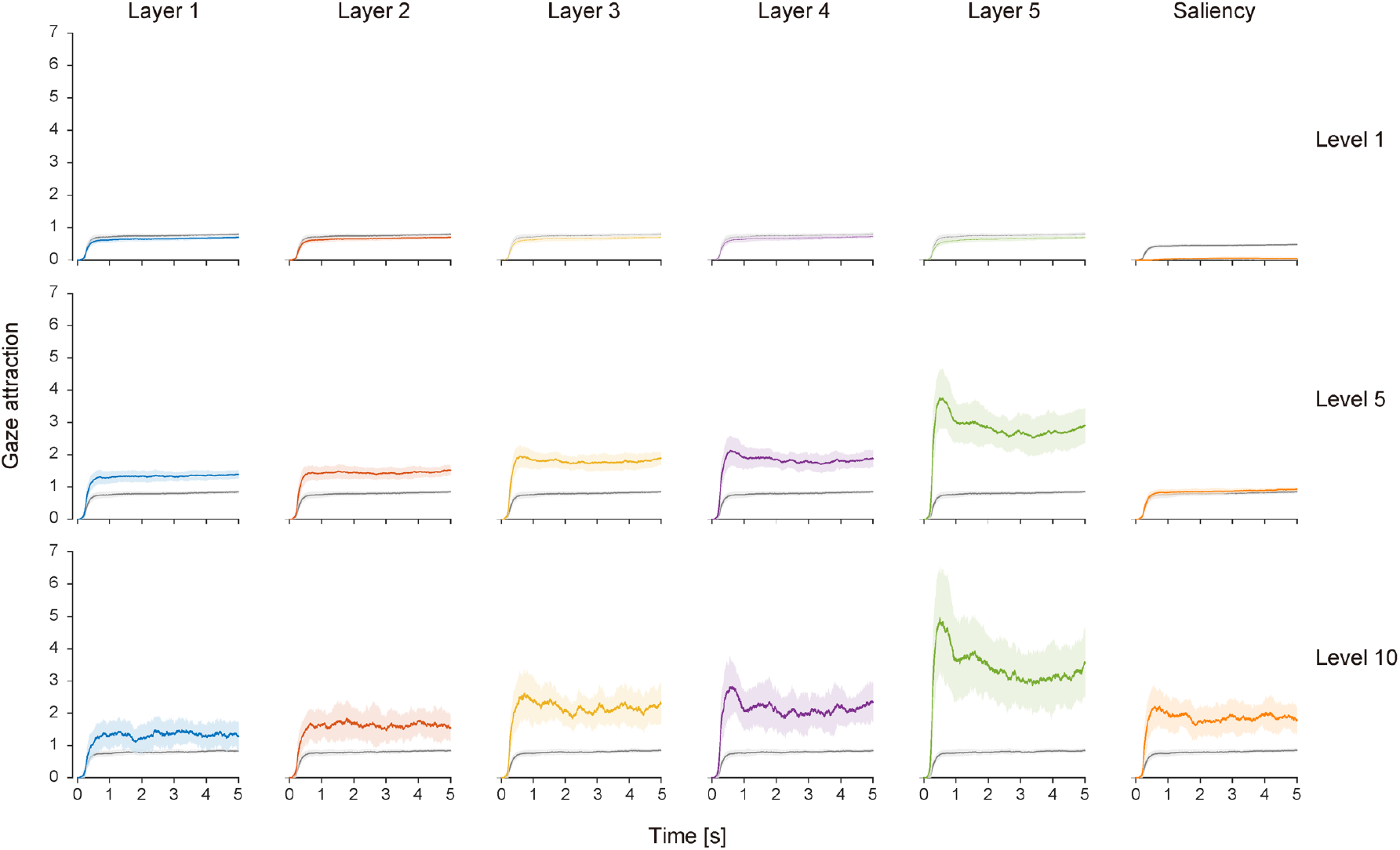
Time courses of gaze attraction for visual features. Three levels of feature intensity (top row, level 1; middle row, level 5; bottom row, level 10) are shown as representative examples. Results from each visual feature are aligned in the column direction. In each graph, the colored solid line and shading indicate the mean and standard deviation over participants, and the gray solid line and shading indicate the mean and standard deviation of the chance level over participants, respectively. The time origin indicates the onset of visual stimulus presentation.

**Figure 4:**
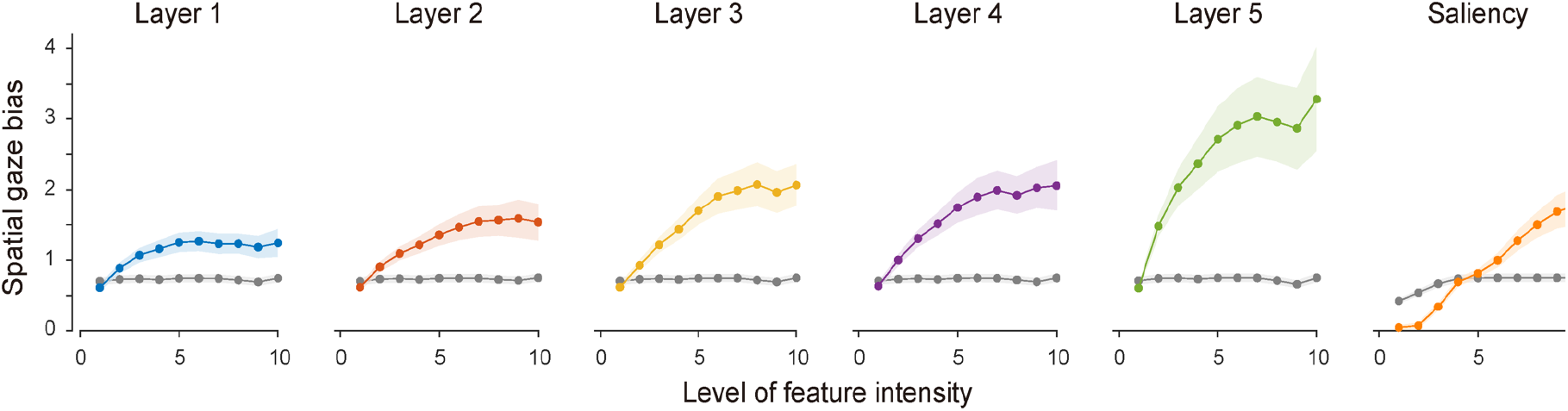
Spatial gaze bias for visual features. The spatial gaze bias for each visual feature is plotted against the feature intensity level. In each graph, the colored solid line and shading indicate the mean and standard deviation over participants, and the gray solid line and shading indicate the mean and standard deviation of the chance level over participants, respectively.

An exceptional result was found for the saliency with lower intensity, where locations with low saliency were seen less than the chance level. We assumed that this result was not due to such locations being avoided but due to the center bias of the gaze (Tatler, 2007) and the artifactual outcomes of saliency computation that output low values near the edge of visual stimuli. More details are given in the Discussion section.

### Temporal gaze bias

The difference between visual features was also evident in terms of the temporal characteristics of the gaze attraction. The time course of the gaze attraction was not flat over time but peaked after the stimulus presentation (at approximately 500 ms), particularly for the visual features corresponding to the deep layers (middle to bottom rows in Figure 3) at higher feature intensity. To quantify this temporal inhomogeneity, we defined the temporal gaze bias, which is a measure of how fast the gaze is attracted to the visual feature (Figure 5A; see details in Methods). Figure 5B shows that the temporal gaze bias was greater for the higher-order visual features activating deep layers of DCNN than the lower-order visual features activating shallow layers of DCNN and saliency, particularly at the higher level of feature intensity. Statistical analyses showed that the temporal gaze bias increased in the following order: saliency, layer 1, 2, 3, 4, and 5 (two-way ANOVA, Bonferroni-corrected p < 0.05 for multiple comparisons; pair-wise significance was not found between saliency vs. layer 1, layer 1 vs. 2, layer 2 vs. 3, layer 3 vs.4; all other pair-wise comparisons were significant). The temporal gaze bias increased as the level of feature intensity except for the shallow layers (p < 0.05 for interaction between visual features and feature intensity, two-way ANOVA), indicating that feature intensity has a strong impact on the gaze attraction in the early period after the stimulus onset particularly for visual features activating the deep layers.

**Figure 5:**
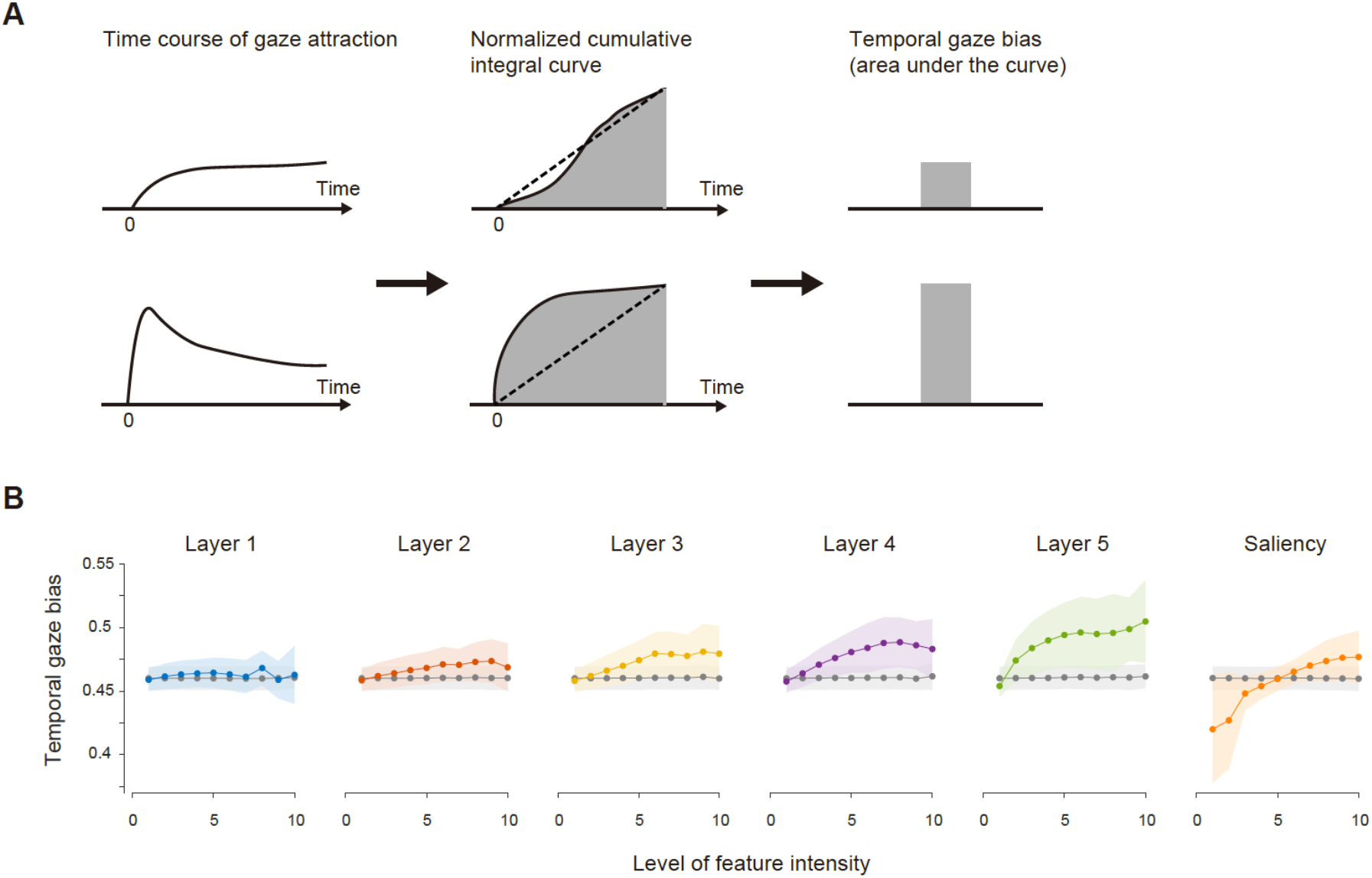
Temporal gaze bias for visual features. A) Procedure in quantifying the temporal gaze bias from the time course of the gaze attraction. The time course of the gaze attraction is integrated cumulatively and then normalized by the maximum value. The convexity of the normalized cumulative integral curve indicates how fast the time course of the gaze attraction reaches a peak value, and the area under the curve serves as a measure of the temporal gaze bias. B) Temporal gaze bias calculated in this procedure. Results are plotted against the feature intensity level. In each graph, the colored solid line and shading indicate the mean and standard deviation over participants, and the gray solid line and shading indicate the mean and standard deviation of the chance level over participants, respectively.

Note that a temporal gaze bias smaller than the chance level was also found for saliency for the same reason described in the previous section (see also the Discussion section).

### Latency of gaze bias

To examine how fast the effect of the gaze attraction emerges, we evaluated the latency of each gaze (up to the 10th gaze) and what visual features attracted the gaze at each gaze order. Figure 6A shows the latency distribution of each gaze order. The first gaze had peak latency at the 260-280 ms bin, which is similar to the result of a previous study using natural scenes as stimuli (Schütt et al., 2019) though a much faster gaze was observed in the distribution. Note that another small peak was found for the first gaze latency at the 80-100 ms bin, but this would be the anticipatory component independent of the visual stimulus observation that remains even after removing such eye movements (see Methods). We then examined what visual feature attracts the first gaze by comparing the gaze attraction of each visual feature. Figure 6B shows that the gaze attraction at the first gaze was significantly higher for the visual feature corresponding to layer 5 than other features (one-way ANOVA for the gaze attraction at the first gaze, Bonferroni-corrected p < 0.05 for multiple comparisons). Figure 6B only shows results at the largest feature intensity level (level 10) as representative examples, but the significance of the layer-5 feature to attract the first gaze was found at level 2 and above (one-way ANOVA for the gaze attraction at the first gaze, Bonferroni-corrected p < 0.05 for multiple comparisons). These results indicate that the deep features attract the human gaze strongly even at the first gaze, and such biases can emerge earlier than 260-280 ms.

**Figure 6:**
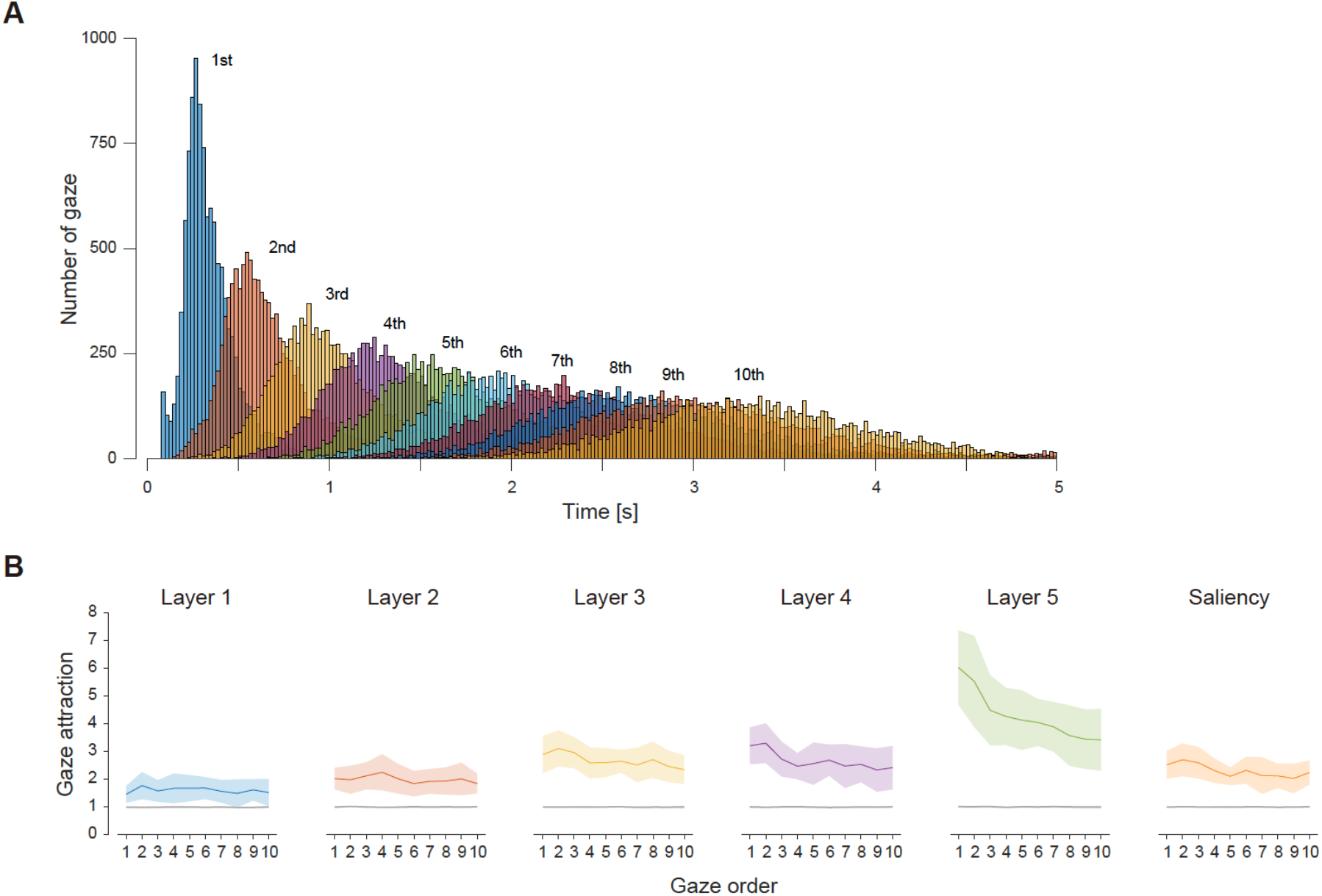
Latency of gaze bias. A) Latency distribution of gaze sorted by the gaze order (up to the 10th gaze; bin width, 20 ms). B) Gaze attraction for visual features sorted by the gaze order. Only results for feature intensity at level 10 are shown here. In each graph, the colored solid line and shading indicate the mean and standard deviation over participants, and the gray solid line and shading indicate the mean and standard deviation of the chance level over participants, respectively.

## Discussion

In this study, we examined what and when visual features are gazed at by measuring eye movements and analyzing visual features embedded in the presented visual stimulus. To reveal the direct relationship between eye movement and visual features, we used a DCNN model by which feature maps can be identified for each visual feature along the DCNN-model hierarchy from the lower (shallow) to the higher (deep) layers. We also used the saliency map for the purpose of comparison as the most conventional visual feature that is considered to attract the gaze. Using the measured gaze data and the feature maps, we quantified the gaze attraction to the visual features by how much they were contained at the gazed location. In contrast with the previous model, our approach did not combine read-out modules with the DCNN model to predict gazed locations but analyzed visual features at the gazed location, allowing us to clarify the direct contribution of each visual feature to the gaze attraction.

As a result, we revealed that higher-order visual features corresponding to the deep layers were a stronger gaze attractor than lower-order visual features corresponding to the shallow layers and the saliency. The spatial locations containing much higher-order visual features were gazed at prominently in the early period after visual stimulus presentation, even at the first gaze, and the preference for the higher-order visual features continued during observation of the visual stimulus. Analyses further showed that this tendency strengthened with the feature intensity, suggesting that the higher-order visual features contribute to the gaze attraction. This is clear experimental evidence of the importance of the higher-order visual features over the lower-order visual features and the conventional saliency model to explain the human gaze bias during the observation of natural scenes.

Then, what areas do play an important role in extracting these higher-order visual features in the human brain? AlexNet, the DCNN model that we used in this study, was developed for image-based object recognition task, and its architecture is designed to follow hierarchical visual systems in the brain, particularly along the ventral pathway. Previous studies showed that neural activity in the higher visual cortex in the ventral pathway of the monkey brain can be predicted from unit activation patterns in the deep layer of AlexNet (Yamins et al., 2014). Human imaging studies also showed that functional magnetic resonance imaging signals of the higher visual cortex in the ventral pathway can be predicted from unit activation patterns in the deep layers of a DCNN model (Güçlü et al., 2015) and vice versa (Horikawa et al., 2017). These studies suggest that the higher visual areas in the ventral pathway are candidates for extracting the higher-order visual features that drive the gaze bias. The strong gaze attraction to a deep feature was observed even at the first gaze, whose peak latency was approximately 260-280 ms. The exact identification of the shortest latency of the first gaze is difficult because of the contamination of anticipatory components, but a reasonable estimate could not be earlier than 120 ms corresponding to the trough of the latency distribution of the first gaze and also roughly corresponding to the shortest end of the second gaze latency distribution (Figure 6A). If we adopt this value as the shortest latency of the gaze bias to the higher-order visual feature, it is slower than the putative activity latency of the higher visual areas of the human brain (Yoshor et al., 2007). It is thus consistent in terms of behavioral and neural latency if the feature extraction in the higher visual areas is the origin of the gaze bias. There are other potential candidates involved in this process including subcortical areas, which could be elucidated in a future study. Combining brain activity measurement with eye movement recording would be suitable for this purpose.

Saliency has been considered a good gaze predictor, particularly in the early period after the stimulus presentation. Such tendency was actually observed in our data. The gaze attraction was greater than chance during the period of visual stimulus presentation, and it had higher values immediately after the visual stimulus presentation, showing that the temporal gaze bias was larger than chance if the feature intensity was sufficient. However, compared with the higher-order visual features, the gaze attraction of the saliency was limited. The gaze attraction to the higher-order visual features was overwhelmingly high in the early period after the visual stimulus presentation, and it continued at a certain level beyond chance until the end of the visual stimulus presentation. These results are consistent with the results of previous studies analyzing the relationship between visual features activating deep layers in a DCNN model and eye movements, though their model was indirect to clarify the contribution of visual features to the gaze bias, suggesting the explanatory power of the higher-order visual features for the spatiotemporal gaze bias.

Another concern about saliency is its multiscale nature based on a Gaussian pyramid (Greenspan et al., 1994), which may complicate the discussion on brain areas involved in its computation. Our results show that the gaze bias of the saliency was approximately between layers 2 and 3 at its highest intensity level, although the saliency computation was based on visual features for which the early visual area has a preference (Koch & Ullman, 1985). There is a discrepancy between brain areas that are supposed to be necessary for the feature preference and the feature extraction scale.

Among the visual features tested in this study, only saliency had a gaze bias lower than chance when the feature intensity was low. This result could be interpreted as participants avoiding directing their gaze at the location with lower saliency. However, it is highly likely that the result is an artifact when computing saliency. The saliency model used in this study outputs a lower value in peripheral areas close to the edges of visual stimuli. Participants usually do not direct their gaze frequently at the peripheral field because of the center bias effect (Tatler, 2007). Therefore, it appears as if participants avoid directing their gaze at the locations with lower saliency, mostly distributed in the peripheral field.

In this study, we found that the higher-order visual features are strong gaze attractors, particularly immediately after the onset of the visual stimulus presentation. This evidence suggests the possibility that a person’s gaze could be guided to specified locations by incepting higher-order visual features with sufficient intensity. Such techniques could be adopted in a critical test of the current findings. The artificial manipulation of visual features could also be useful in designing traffic signs, fail-free user interfaces, and effective advertisements. Thus, our approach could provide a new method of exploring and even utilizing the relationship between visual features and induced eye movement.

## Methods

### Participants

Twenty participants (16 males, 4 females; age ranging between 20 and 26 years (mean, 21.7 years)) participated in the experiment. All participants had normal or corrected-to-normal vision acuity. They were compensated by 1000 JPY per hour for their participation. All participants gave written informed consent before participating in the experiments. The procedure was approved by the institutional review board of the University of Electro-Communications.

### Visual stimuli

Images of natural scenes were selected from the ADE20K database (Zhou et al., 2017) and Pascal Context database (Mottaghi et al., 2014), in which objects are segmented separately and annotated with corresponding object categories. We used color images of more than four object categories with a (horizontal-to-vertical) aspect ratio of 4 to 3 as visual stimuli for the experiment. The number of selected images was 496 from ADE20K and 154 from Pascal Context, respectively. Each image was rescaled to 800 ×600 pixels (width ×height).

### Apparatus

Visual stimuli were presented on a 21-inch CRT monitor (FlexScan T966, EIZO NANAO Inc.; frame rate, 60 Hz; resolution, 1024 ×768 pixels (width ×height)). Participants sat 86 cm from the monitor, and the visual stimuli subtended a field of view of approximately 20 deg ×15 deg (width ×height). A chin rest was used to keep the participant’s head still. During the visual stimulus presentation, the participant’s right eye movements were recorded with an EyeLink 1000 (SR Research Ltd.) desktop mount system at a sampling rate of 1000 Hz. An image of a small white square was displayed synchronously with the visual stimuli at the left side of the CRT monitor, and a light sensor was attached on the monitor to detect the onset of visual stimulus presentation. The detected onset was used to precisely define the time origin for the recoded eye movement. The light sensor was covered with a dark cloth so that participants saw neither the white square nor the light sensor during the experiment. Visual stimulus presentation was controlled by MATLAB (The MathWorks, Natick, MA) using Psychophysics Toolbox Version 3 (Pelli, 1997; Brainard, 1997; Kleine et al., 2007).

### Experimental design

The experiments comprised encoding and recognition runs. In the encoding runs, the participants were asked to observe presented visual stimuli with eye movements allowed. In the recognition runs, the participants were asked to observe presented visual stimuli with eye movements allowed and to answer whether each stimulus had been presented in the preceding encoding runs.

After the eye tracker was calibrated using a nine-point fixation presented on the monitor, three sets of the combination of an encoding run and a recognition run were performed (Figure 1). There were 40, 39, and 39 stimulus presentation trials in the encoding runs, each of which was followed by eight stimulus presentation trials in the recognition runs. This sequence (calibration and the three sets of the combination of the encoding and recognition runs) was repeated five times, resulting in 590 (450 from ADE2K and 140 from Pascal Context) and 120 (92 from ADE2K and 28 from Pascal Context, with half being the same images as those in the encoding runs for the recognition task) natural scene presentations for encoding and recognition, respectively. The eye tracker was recalibrated between runs when necessary.

In the encoding runs, the participants pressed the space key on a keyboard to start each trial. A white fixation cross (0.3 deg ×0.3 deg) was then presented for 500 ms at the center of the CRT monitor on a black background, and an image of a natural scene was subsequently presented as a visual stimulus. The participant was instructed to observe the presented stimulus with free eye movements while keeping their head still on the chin rest. The presented stimulus automatically disappeared at 5000 ms after the onset of the stimulus presentation, and the monitor turned to black until the participant pressed the space key to start the next trial (the instruction to press a button appeared on the screen; see Figure 1A). The participant’ right eye movements were continuously monitored during the trial. The eye tracker automatically recognized saccade events if the speed of eye movement exceeded 30 deg/s, and the remaining events were recognized as fixation events after removing blinks. The spatial coordinates and event time of each fixation event recorded by the eye tracker provided information on the gazed spatial location and gaze time after the onset of stimulus presentation. Fixation events that happened before 80 ms after stimulus presentation were considered anticipatory and excluded from the analysis (Gezeck et al., 1997).

In the recognition runs, the participants again pressed the space key on a keyboard to start each trial. A white fixation cross (0.3 deg ×0.3 deg) was presented for 500 ms at the center of the CRT monitor on a black background, and an image of a natural scene that was either one presented in the preceding encoding run or one new to the experiment was then presented as a visual stimulus. There were four previously presented stimuli in each recognition run; i.e., the probability of seeing a previously presented image was 50%. The participant was instructed to observe the visual stimuli with free eye movements while keeping their head still on the chin rest. The presented stimuli automatically disappeared at 2000 ms after the onset of the stimulus presentation and the monitor turned black. The participant was instructed to press the “z” key on the keyboard during this period if they thought the stimulus had been presented in the preceding encoding run or press the slash key if not, without a time constraint (the instruction to press a button appeared on the screen; see Figure 1A). After answering the task, the participants pressed the space key to start the next trial.

### Visual feature analysis

#### DCNN feature map

DCNN feature maps for each image were computed using AlexNet pretrained with the ImageNet database (Deng et al., 2009) to classify images into 1000 object categories. AlexNet comprised five convolutional layers and three fully connected layers. In this study, we generated feature maps from the five convolutional layers using SmoothGrad (Smilkov et al., 2017). SmoothGrad can generate a map that shows the intensity distribution of visual features activating a specified layer of the DCNN by backpropagating all unit activations in the layer into the image-pixel space. This procedure was repeated for all images and for all five convolutional layers of the DCNN, resulting in five DCNN-feature maps (from layers 1 to 5) for each image (Figure 2A).

#### Saliency map

A saliency map for each image was computed using SaliencyToolbox (Walther and Koch, 2006), a typical saliency model that combines conspicuity maps of luminance contrast, color contrast, and orientation. Here, these maps were combined with equal weights to compute saliency. The saliency map shows a distribution of the intensity of saliency, similarly to the DCNN-feature maps (Figure 2A).

### Quantitative evaluation of gaze attraction to visual features

In this study, we defined the gaze attraction to a particular visual feature by how much the visual feature is contained at the spatial location where the gaze was attracted (Figure 2B). To consider the effect of the intensity difference in visual features, we first rescaled visual feature intensity into the range between 0 (minimum) and 1 (maximum) for each image and discretized it into ten levels. Given the visual feature maps labeled with the feature intensity levels, we could identify what visual features were dominant at the gazed location and thus derive a quantity for the gaze attraction. However, this procedure might be inaccurate because the gazed locations identified by the eye tracker contain measurement errors, which are expected to be within 1 deg of the visual angle according to the specifications of the eye tracker. Therefore, instead of using the feature dominance at a single point in the image, we defined a circular region of 1-deg radius around each gazed location and calculated how much of the area was occupied by each visual feature (different intensity levels were treated separately) in the region. If there is a difference in the total area occupied by each visual feature in the entire image, it yields spurious biases in the occupancy rate of each visual feature in the gazed region. To avoid such effects, we divided the occupancy rate of each visual feature in the gazed region by the total occupancy rate of each visual feature in the entire image. This relative occupancy rate thus indicates how dominant each visual feature is specifically in the gazed region, which we define as the gaze attraction of the visual feature. Repeating this procedure for each gazed location at its gaze time and taking the average over all presented stimuli (Figure 2B), we obtained the time course of the gaze attraction for each visual feature at different intensity levels (Figure 3).

#### Spatial gaze bias

To examine spatial locations that attract the human gaze, conventional studies typically calculated a fixation map that represented the frequency with which the gaze was directed at each location during visual stimulus presentation. This reflects the gaze bias to each location in presented visual stimuli because all the gaze is counted for the duration of visual stimulus presentation while the temporal difference in the gaze frequency is ignored. Similarly, to evaluate how frequently the gaze was attracted to spatial locations containing each visual feature with a specified intensity level, we simply took the average of the time course of the gaze attraction for each visual feature at the intensity level. We defined this quantify as the spatial gaze bias to the visual features.

#### Temporal gaze bias

The gaze attraction is not constant but changes dynamically over time. As shown in Figure 3, some visual features had a clear peak in the early period after visual stimulus presentation and others did not. To quantify such a difference in temporal characteristics, we took a cumulative integral over the time course of the gaze attraction (Figure 5A). The time course of the gaze attraction comprises non-negative values, and its cumulative integral thus becomes convex if the time course shows higher values close to the onset of stimulus presentation whereas it becomes concave if the time course shows the opposite tendency. Thus, the convexity of the cumulative integral of the time course of the gaze attraction reflects how quickly the visual feature is gazed at after visual stimulus presentation. We here quantified the convexity using the area under the cumulative integral of the time course of the gaze attraction. We defined this quantity as the temporal gaze bias to the visual features. Note that the time course of the gaze attraction was normalized such that its minimum-to-maximum range becomes 0 to 1 before computing the temporal gaze bias because the constant component of the time course affects the convexity of the cumulative integral although it is irrelevant to the temporal gaze bias. The cumulative integral was also normalized such that its minimum-to-maximum range became 0 to 1 so as to evaluate its convexity independently of the absolute values of the cumulative integral.

We confirmed that this procedure faithfully measures the temporal gaze bias by simulation analysis using hypothetical time courses of the gaze attraction (see Supplementary information). The temporal gaze bias varied consistently with model parameters that control the peak time (Supplementary figure 1A, 1B) and the magnitude of the transient component (Supplementary figure 1C, 1D) of the gaze attraction.

#### Chance levels of spatial and temporal gaze bias

It is difficult to define the chance levels of the spatial and temporal gaze bias theoretically, and we thus adopted the following empirical procedure to define them. For each participant, gaze coordinates measured for all 590 trials in the encoding runs were discarded and random coordinates uniformly sampled from the entire image dimension were reassigned for all gaze data while the timings were kept unchanged. The time course of the gaze attraction was then computed in the same way as for the original gaze data. This operation was conducted 100 times and the time courses were averaged. The averaged time course of the gaze attraction with randomized coordinates keeps the mean frequency of the gaze at each timing (because the timing information is preserved), but the occupancy rate of each visual feature in a reassigned gazed location changes and asymptotically approaches the mean density of each visual feature in the entire image, serving as the time course of the chance-level gaze attraction for each visual feature for each participant. The spatial and temporal gaze bias was calculated from the time course of the chance-level gaze attraction using the same procedure explained above.

## Acknowledgment

We thank Edanz (https://jp.edanz.com/ac) for helping edit a draft of this manuscript. This research was supported by JST PRESTO Grant Number JPMJPR1778, JSPS KAKENHI Grant Number 20H00600, 18KK0311, and 17H01755, and Yazaki Memorial Foundation for Science and Technology.

## Author contributions

KA and YM developed the concept; KA and YM developed the experiment design and protocol; KA collected and analyzed the data; KA and TN developed methods for image analyses; KA and YM wrote the paper. All authors reviewed the final manuscript.

