## Supplementary information for "Spatiotemporal bias of the human gaze toward hierarchical visual features during natural scene viewing"

### Validation of temporal gaze bias using a mathematical model

To validate the temporal gaze bias defined in this paper, we used a mathematical equation to model the time course of the gaze attraction whose peak timing varies, and examined whether the temporal gaze bias can capture those temporal variations.

The time course of the gaze attraction, $f(t)$, was modeled as a Gaussian function for simplicity,

$$f(t) = \exp\left[-\frac{(t-\mu)^2}{2\sigma^2}\right], \quad (1)$$

where $t$ is the elapsed time after the onset of stimulus presentation, $\mu$ and $\sigma$ are the peak time and the width of the gaze attraction, respectively. Supplementary figure 1A shows the modeled time course of the gaze attraction for three different values of $\mu$ ($\sigma$ was fixed at 0.175 s for all cases). Supplementary figure 1B shows the temporal gaze bias calculated by the same method described in Methods section in the main text of the paper. The temporal gaze bias gradually decreased as the peak time, demonstrating that our metric faithfully represents whether the gaze was biased to the early period within a trial.

Although this model is an easy example to understand how the temporal gaze bias changes according to the peak time, observed time courses did not have a shape like them but rather looked like a combination of a transient component and a sustained component (see Figure 3 in the main text). To represent these components, we tested a modified model comprising a Gaussian transient component and a sigmoid sustained component

$$g(t) = A_t \exp\left[-\frac{(t-\mu_t)^2}{2{\sigma_t}^2}\right] + A_s \frac{1}{1+\exp\left[-c(t-\mu_s)\right]}, \quad (2)$$

where $\mu_t$ and $\sigma_t$ are the peak time and the width of the transient component, $\mu_s$ and $c$ are the temporal shift and the ramp-up gain of the sustained component, and $A_t$ and $A_s$ are the amplitude of the transient component and sustained component, respectively. Supplementary figure 1C shows four cases modeling the time course of the gaze attraction synthesized using different combinations of $A_t$ and $A_s$ while other parameters were fixed ($\mu_t$, 0.5 s; $\sigma_t$, 0.175 s; $\mu_s$, 0.25 s; $c$, 20 /s). The case 1 and 2 had the transient component with different amplitudes (the case 1 was twice as large as the case 2) and the sustained component whose amplitude was the same as the transient component of the case 2. The case 3 and 4 had only the sustained component. The amplitude of the sustained component of the case 3 was the same as that of the case 2. The amplitude of the sustained component of the case 4 was set so that the maximum value matched that of the case 1. Supplementary figure 1D shows the temporal gaze bias for these time courses. The temporal gaze bias decreased when the amplitude of the transient component decreased (case 1 to 2) but was higher than the case 3 and 4 that had no transient component. Note that the temporal gaze bias of the case 4 was smaller than that of the case 1 and 2, and it was actually the same as that of the case 3 even though the time course of the case 4 had a maximum value higher than that of the case 3. These results prove that the temporal gaze bias correctly captures the existence of the transient component irrespectively of the overall magnitude of the time course of the gaze attraction, indicating that our metric is valid to quantify whether the gaze attraction has large values in the early period.

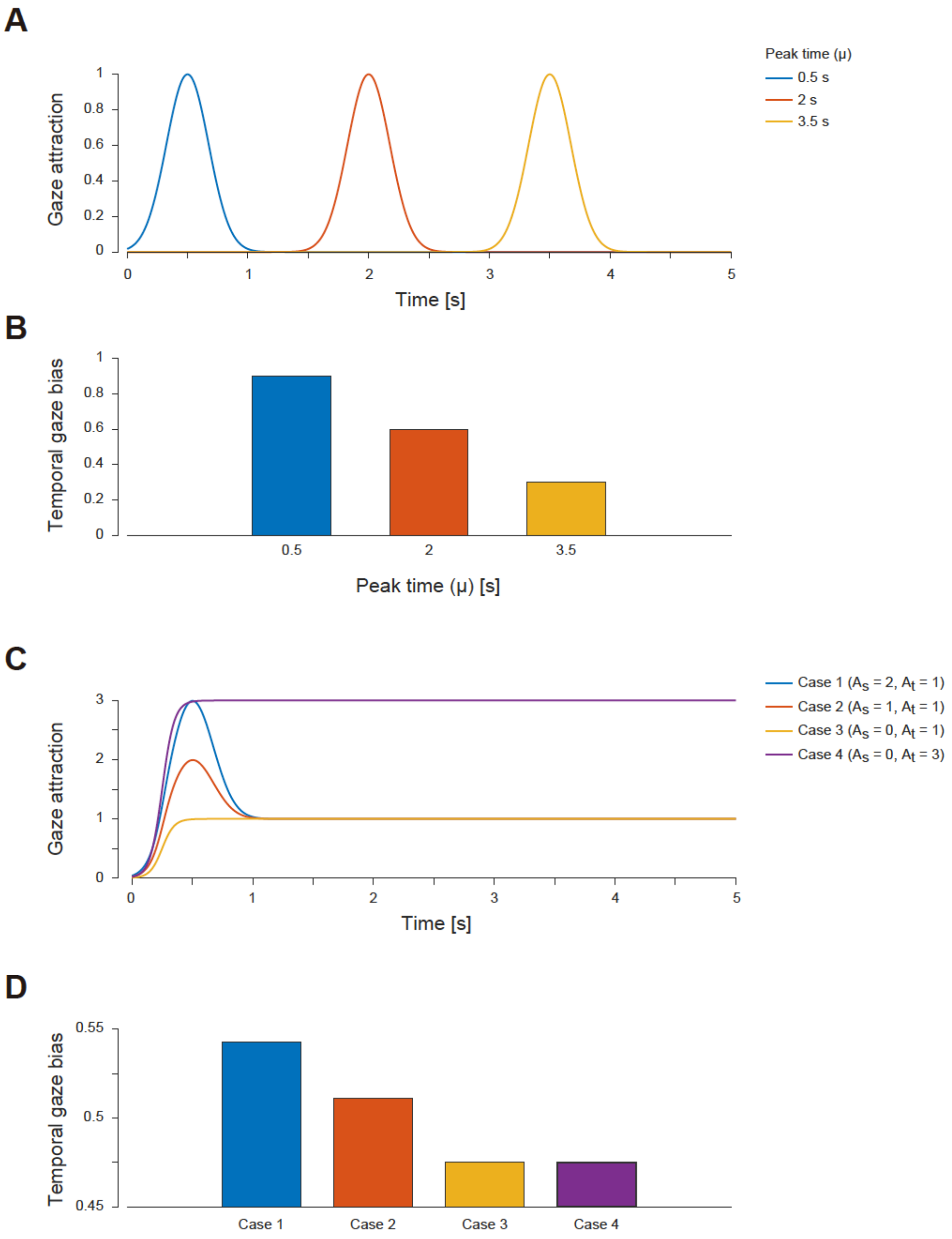


**Supplementary figure 1:** Temporal gaze bias for the time course of the gaze attraction synthesized by a mathematical model. A) Examples of the model time courses of the gaze attraction with different peak times. B) Temporal gaze bias calculated for the time courses of the gaze attraction presented in panel A. C) Examples of the model time courses of the gaze attraction comprising the transient and the sustained component. The mixing coefficients were varied as displayed in the legend. Other parameters of the model were identical. D) Temporal gaze bias calculated for the time courses of the gaze attraction presented in panel C.

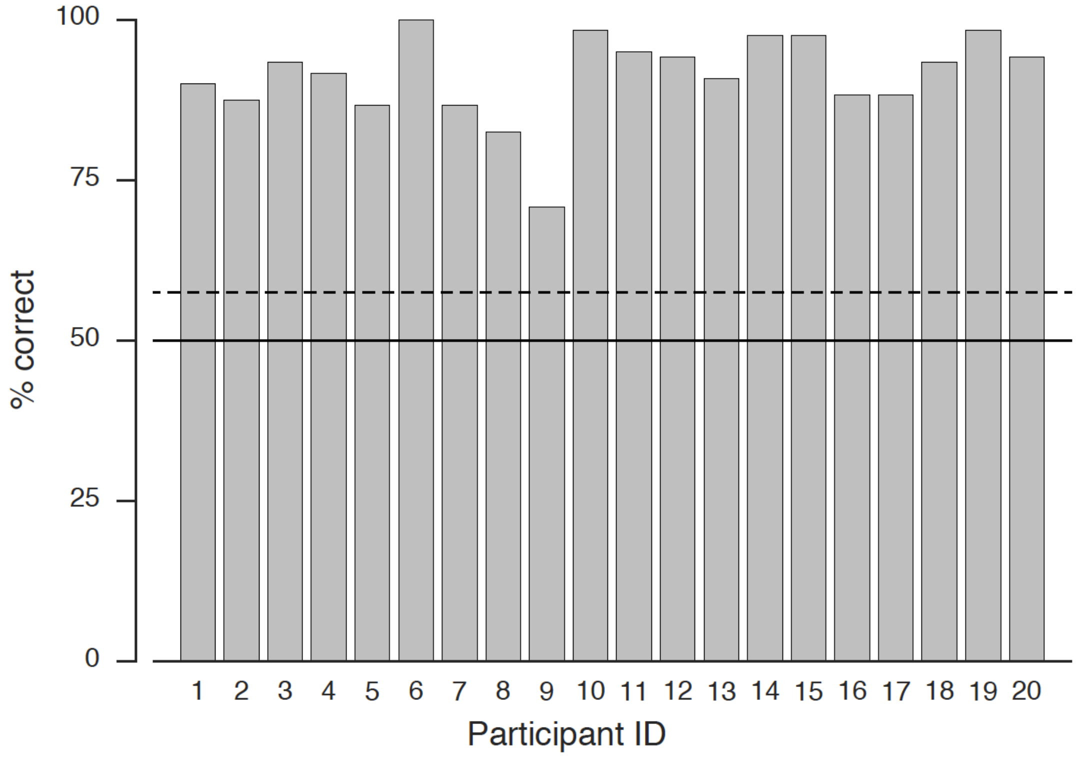


**Supplementary figure 2:** The recognition task performance. The percentage of correct of the recognition task was significantly higher than the chance level (57.5% represented by the dashed line, $p < 0.05$ for binomial test) for all participants. The mean of the percentage of correct over participants was 91.25% and the standard deviation was 6.74%.
